# Identifying Pathogen and Allele Type Simultaneously (IPATS) in a single well using droplet digital PCR

**DOI:** 10.1101/2022.09.10.507438

**Authors:** Kosuke Notsu, Hala El Daous, Shuya Mitoma, Xinyue Wu, Junzo Norimine, Satoshi Sekiguchi

**Affiliations:** Graduate School of Medicine and Veterinary Medicine, University of Miyazaki, Miyazaki 889-1692, Japan; Faculty of Veterinary Medicine, Benha University, Toukh 13736, Egypt; Department of Veterinary Science, Faculty of Agriculture, University of Miyazaki, Miyazaki 889-2192, Japan; Center for Animal Disease Control, University of Miyazaki, Miyazaki 889-2192, Japan

**Author notes:** **Corresponding author**: Satoshi Sekiguchi, Department of Veterinary Sciences, Faculty of Agriculture, University of Miyazaki, 1-1, Gakuen-Kibanadai-Nishi, Miyazaki 889-2192, Japan.

## Abstract

A combined host biomarker and pathogen diagnosis provides insight into disease progression risk and contributes to appropriate clinical decision-making regarding prevention and treatment. In preventive veterinary medicine, such combined diagnosis could improve risk-based livestock herd management. We developed a single-well based test for combined diagnosis of bovine leukemia virus (BLV) and bovine MHC (*BoLA*)-*DRB3* alleles. A fourplex droplet digital PCR method targeting the BLV *pol* gene, BLV-susceptible *DRB3*016:01* allele, resistant *DRB3*009:02* allele, and housekeeping RPP30 gene (IPATS-BLV) successfully measured the percentage of BLV-infected cells and determined allele types precisely. Furthermore, it discriminated homozygous from heterozygous carriers. Using this method to determine the impact of carrying these alleles on the BLV proviral load (PVL), we found *DRB3*009:02-carrying* cattle could suppress the PVL to a low or undetectable level, even with the presence of a susceptible allele. Although the population of *DRB3*016:01*-carrying cattle showed significantly higher PVLs when compared with cattle carrying other alleles, their individual PVLs were highly variable. Because of the simplicity and speed of this single-well assay, IPATS could be a suitable platform for the combined diagnosis of host biomarkers and pathogens in a wide range of other systems.

## Background

Owing to decades of effort associating genetic information with disease risk, genomic risk prediction is now being implemented clinically (Abraham & Inouye, 2015; Lewis & Vassos, 2020). It contributes to clinician decision-making regarding disease treatment and prevention, and it provides more flexible customized treatment for patients. Overcoming the threat of infectious disease requires an accurate risk prediction of the disease severity for individuals. However, even among those harboring disease-related biomarkers, many of which have been identified in population level studies, disease progression and its outcome vary because of the highly complex and dynamic host–pathogen interactions that occur during infection (Ellner et al., 2021; Zhang et al., 2022). A combined diagnosis that encompasses genomic risk prediction and pathogen identification is one potential solution for overcoming the heterogeneity of individual infection and would provide deeper insight into an individual’s infectious status.

Human leukocyte antigen (HLA; major histocompatibility complex (MHC) in humans), proteins on the surface of cells are involved with the regulation of innate immunity and antigen presentation (Neefjes et al., 2011; Wieczorek et al., 2017). The HLA haplotype is informative for predicting the strength of an individual’s immune responses against pathogens and is a useful indicator of disease susceptibility (Augusto & Hollenbach, 2022; Matzaraki et al., 2017). The impact of differing immune capacity against viral replication could result in the emergence of a rare population with highly transmissibility to others (super spreaders). Additionally, high immune capacity can suppress an individual’s viral load to levels undetectable by diagnostic testing (e.g., elite controllers in human immunodeficiency virus studies). In viral infections, the viral load is diagnostically important because it acts as an indicator of disease severity (Fajnzylber et al., 2020; Granados et al., 2017; Yamano et al., 2002) and transmissibility (Attia et al., 2009; Marc et al., 2021; Marks et al., 2021). Determining both the HLA haplotype and viral load has the benefit of accurately identifying infection susceptible/severe disease patients and infection resistant/mild disease patients. Such information supports the prioritization of intensive medicine and vaccination to the at-risk population.

There is a critical need of diagnostics that determine the genomic risk prediction and quantity of pathogens for livestock infectious diseases. The eradication of highly contagious diseases, such as foot and mouth diseases (Knight-Jones & Rushton, 2013) and African swine fever (Mason-D’Croz et al., 2020), and of chronic, untreatable diseases, such as paratuberculosis (Johne’s disease) (Garcia & Shalloo, 2015) and bovine leukemia virus (BLV) infection (Pelzer, 1997), is an unavoidable challenge to assure future food production. These diseases are listed as notifiable terrestrial animal diseases by the World Organization for Animal Health (World Organization of Animal Health, 2022). The latter diseases are difficult to control because of their silent spread, owing to the lack of clinical signs, and the unfeasibility of culling of all infected animals, owing to the high prevalence worldwide (Polat et al., 2017; Whittington et al., 2019). To control these diseases while preserving as many animals as possible, identifying and isolating susceptible super spreader animals and maintaining disease-resistant animals via selective breeding is a reasonable approach.

BLV belongs to the genus *Deltaretrovirus* in the Retroviridae family, and it has a genomic structure and properties similar to those of human T-lymphotropic virus type 1 (Aida et al., 2013). BLV causes production issues in livestock farms by reducing the milk and meat productivity of infected cattle (Nakada et al., 2022; Ott et al., 2003). Furthermore, just under 10% of BLV-infected cattle develop a malignant B-cell lymphoma called enzootic bovine leukosis (EBL) upon lifelong infection (Burny et al., 1988). As there are no effective treatments or vaccines for BLV infection (Barez et al., 2015), an appropriate intervention to prevent the spread of this virus is needed. BLV transmits via the direct transfer of infected blood, so the proviral load (PVL) is a determinant of transmissibility. Previous research revealed an association between exon 2 of the bovine MHC (*BoLA*)-*DRB3* gene (*DRB3*) and the BLV PVL. In the Japanese Black species of cattle, having *DRB3*016:01* is associated with a high PVL (HPL) of BLV; thus, this allele is considered to be a BLV susceptibility gene (Lo et al., 2021; Miyasaka et al., 2013). In contrast, having *DRB3*009:02* is strongly associated with a low PVL (LPL) of BLV in the Japanese Black and Holstein species of cattle; thus, this allele is considered to be a BLV resistance gene (Carignano et al., 2017; El Daous et al., 2021; Hayashi et al., 2017; M. A. Juliarena et al., 2008; Lo et al., 2021; Miyasaka et al., 2013; Takeshima et al., 2019). To provide a method for identifying BLV-susceptible super spreaders and BLV-resistant elite controllers more easily and rapidly, we developed a single-well droplet digital PCR (ddPCR)-based measurement system for the BLV PVL, *DRB3*016:01* allele, and *DRB3*009:02* allele.

## Results

### *Single-well measurement of BLV PVL, DRB3*016:01,* and *DRB3*009:02*

This study aimed to design a method for easily and rapidly identifying BLV-susceptible super spreaders (*DRB3*016:01*-carrying cattle with a HPL of BLV) and BLV-resistant elite controllers (*DRB3*009:02*-carrying cattle with an undetectable PVL). We developed a fourplex ddPCR targeting the BLV *pol* gene, *DRB3*016:01* allele, *DRB3*009:02* allele, and housekeeping gene RPP30, named IPATS (<u>I</u>dentifying <u>P</u>athogen and <u>A</u>llele <u>T</u>ype <u>S</u>imultaneously)-BLV (Figure 1A–1H). This assay consists of a multiplex TaqMan assay using seven primers, including two locked nucleic acid (LNA) primers, and four TaqMan probes in a single well (Tables S1 and S2). By modulating the amplicon length and primer/probe concentration in the reaction mixture, we succeeded at detecting two targets in the same color with separate fluorescence magnitudes in the PCR-positive droplet (Levy et al., 2021; Miotke et al., 2014). When a droplet contains a *DRB3*016:01* allele or/and *DRB3*009:02* allele, which we set to be detected by a low-concentration probe (approximately 200 bp of amplicon), the droplet exhibits a low level of FAM or/and HEX color, respectively, in our TaqMan assay. When a droplet contains a BLV *pol* gene or/and RPP30, which we set to be detected by a high-concentration probe (approximately 100 bp of amplicon), the droplet exhibits a high level of FAM or/and HEX color, respectively, in our TaqMan assay. When a droplet contains both *DRB3*016:01* and the BLV *pol* gene (i.e., low and high levels of FAM color) or *DRB3*009:02* and RPP30 (i.e., low and high levels of HEX color), a cluster showing a very high level of color is observed (Figure 1D–1F). This assay visualizes the properties of BLV PVL, *DRB3*016:01* allele presence, and *DRB3*009:02* allele presence in samples via the FAM and HEX amplitude cluster patterns of droplets (Figure 1F). We used the percentage of BLV-infected cells as an indicator of the BLV PVL. We could calculate the percentage of BLV-infected cells by dividing the number of BLV-positive droplets by half of the number of RPP30-positive droplets. Furthermore, this assay can determine the heterozygosity or homogeneity of *DRB3*016:01* and *DRB3*009:02* by dividing the number of *DRB3*016:01*/*\*009:0*2-positive droplets by the number of RPP30-positive droplets.

**Figure 1.**
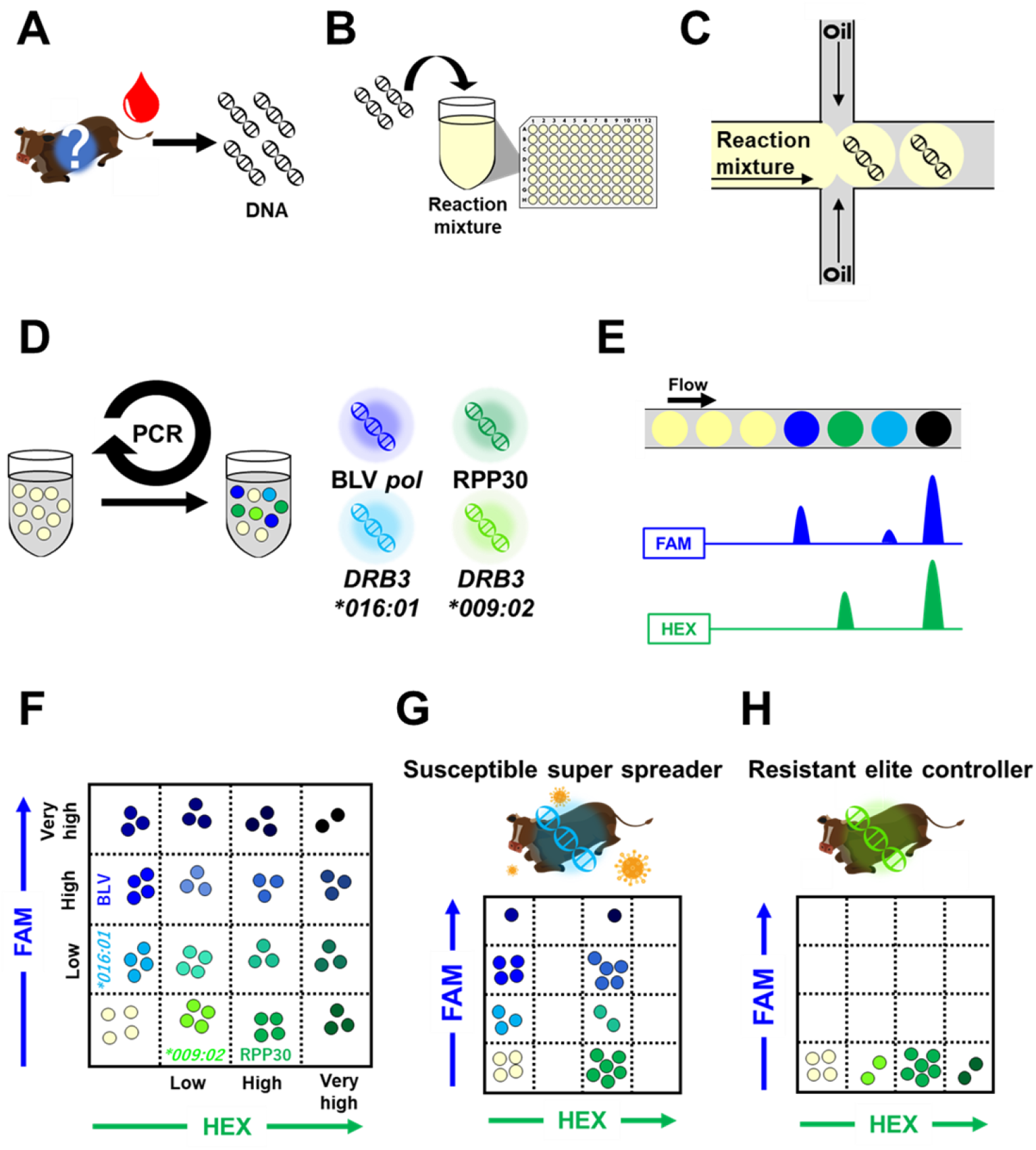
Workflow of IPATS-BLV. The work flow is indicated from a to f. (A) DNA extraction from bovine whole blood. (B) Addition of DNA samples to the reaction mixture. (C) Generation of droplets for partitioning the sample DNA. (D) Fourplex TaqMan Assay of the droplets. (E) Determination of the florescence magnitude. (F) 2D amplitude indicating the position of droplet clusters according to the fluorescence magnitude. (G) 2D amplitude pattern of a BLV-susceptible super spreader. (H) 2D amplitude pattern of a BLV-resistant elite controller.

As shown in Figure 2A–2I, IPATS-BLV produces a variety of cluster patterns of FAM and HEX fluorescence intensity in 2D amplitude. BLV-susceptible super spreaders produce the cluster patterns shown in Figure 2D and 2C (Figure 2C displays the pattern produced by *DRB3*016:01/*009:02*-carrying cattle with a HPL of BLV). In contrast, BLV-resistant elite controllers produce the cluster patterns shown in Figure 2B and 2D (Figure 2D displays the pattern produced by *DRB3*016:01*/* *009:02*-carrying cattle with an undetectable PLV of BLV). *DRB3*016:01-carrying* cattle with an undetectable PVL of BLV produce the pattern shown in Figure 2E. When cattle carry neither the *DRB3*016:01* allele nor the *DRB3* 009:02* allele, those with a HPL, LPL, or undetectable PVL of BLV produce the cluster patterns shown in Figure 2F, 2G, and 2H, respectively. Figure 2I displays the pattern produced by cattle that are negative for all the target genes. A 1D amplitude of these patterns is provided in Figure S1A-S1I.

**Figure 2.**
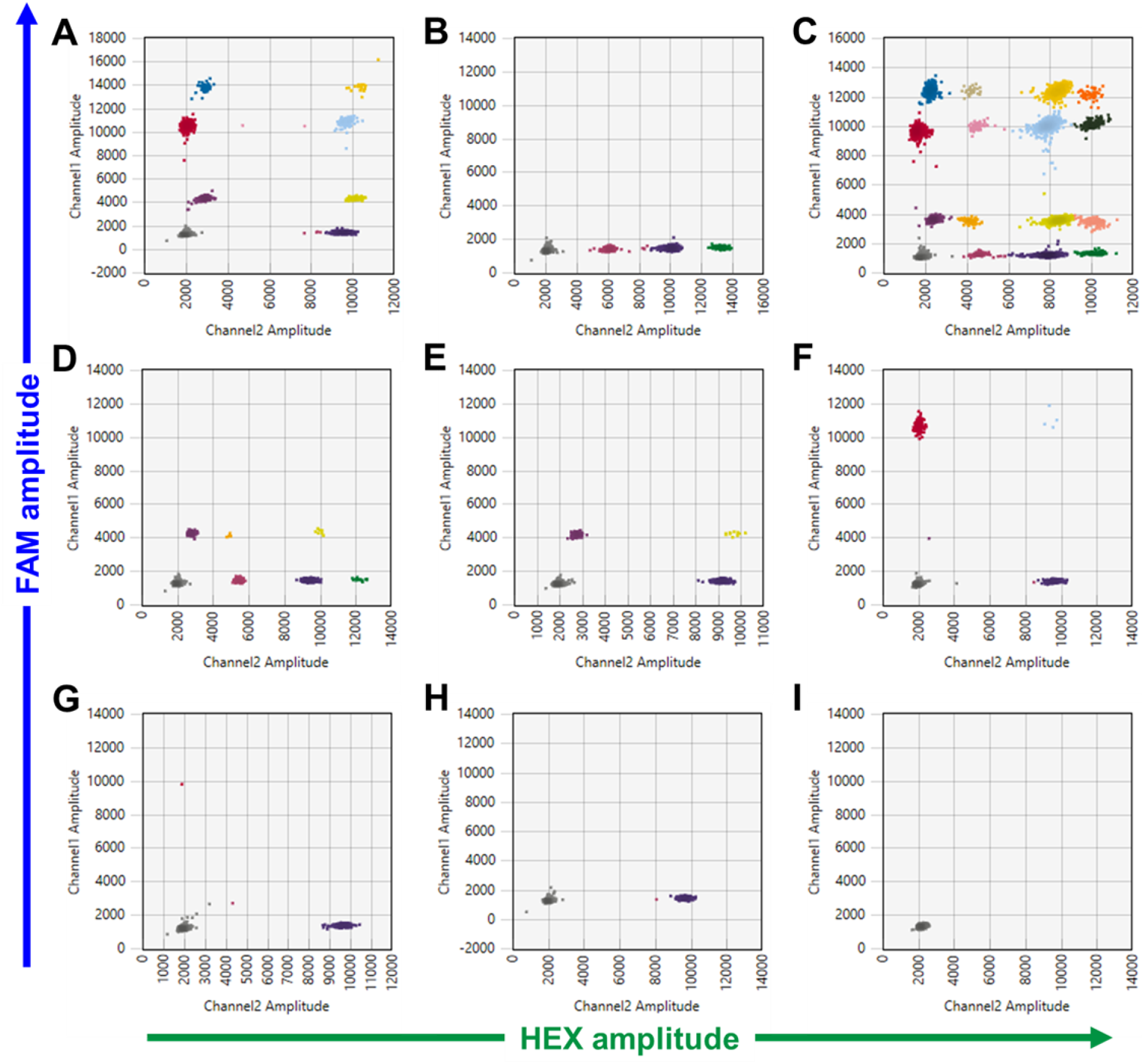
Cluster patterns in IPATS-BLV 2D amplitudes. Cluster patterns of IPATS-BLV of eight cattle with different possession of *DRB3*016:01, DRB3*009:02* and BLV provirus, and water is shown. Each droplet produces each different FAM and HEX fluorescence magnitude in TaqMan assay, reflecting a presence of targeting genes within droplet. Droplets makes clusters according to the similarity of fluorescence magnitude. The divisions of clusters are indicated by different color of droplets. (A) BLV-susceptible super spreader. (B) BLV-resistant elite controller. (C) Mixed population of *DRB3*016:01/*015:01-carrying* cattle with a HPL of BLV and *DRB3*009:02/*015:01*-carrying cattle (presumably *DRB3*009:02/*016:01* heterozygous with detectable BLV provirus). (D) *DRB3*009:02*/*\*016:01* heterozygous cattle with undetectable BLV provirus. (E) *DRB3*016:01*-carrying cattle with undetectable BLV provirus. (F) Other allele-carrying cattle with a HPL of BLV. (G) Other allele-carrying cattle with a LPL of BLV. (H) Other allele-carrying cattle with undetectable BLV provirus. (I) Water.

### Digital allele typing with high accuracy

To assess the accuracy of the *DRB3*016:01* and *DRB3*009:02* genotyping by our new method, we performed IPATS-BLV on 58 bovine genomic DNA samples with *DRB3* allele variation. These samples were previously genotyped using combined PCR-Restriction Fragment Length Polymorphism (RFLP)-sequencing (El Daous et al., 2021; Notsu et al., 2022; Van Eijk et al., 1992). A total of 21 *DRB3* alleles were identified by this previous analysis (Table S3). Among these samples, IPATS-BLV successfully discriminated seven samples with *DRB3*016:01* alleles and 14 samples with *DRB3*009:02* alleles by calculating the ratio of the number of *DRB3*016:01-positive (DRB3*016:01* ratio) and *DRB3*009:02*-positive (*DRB3*009:02* ratio) droplets to the number of RPP30-positive droplets (Table 1 - Table S3). Five of these samples were *DRB3*016:01/*009:02* heterozygous. The *DRB3*016:01* and *DRB3*009:02* ratio was 0.4646 (standard error (SE): ±0.01087) and 0.4658 (SE: ±0.00779), respectively. Because we rounded the values of the *DRB3*016:01* and *DRB3*009:02* ratios of samples carrying other alleles to two decimal places to suppress the effect of noise, all these samples had values of 0.0 for their ratios, except for a sample carrying heterozygous *DRB3*037:01/*044:01* (yellow-highlighted in Table S3) which had a *DRB3*016:01* ratio value of 0.2. Thus, IPATS-BLV had a 100% diagnostic sensitivity and specificity for *DRB3*016:01* and *DRB3*\**009:02* genotyping.

**Table 1.**
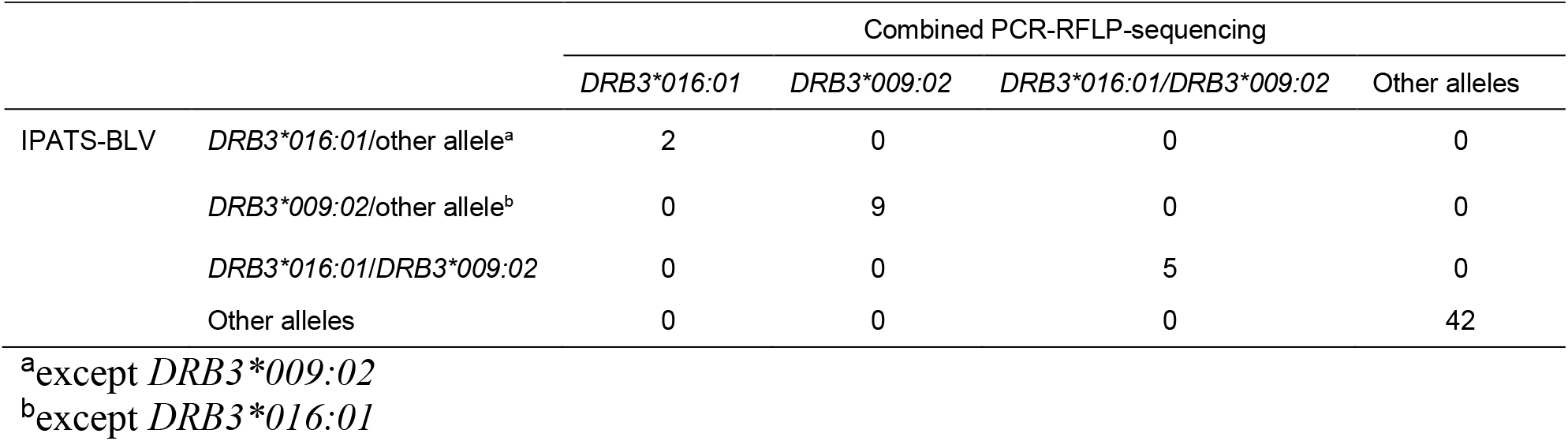
Comparison of the allele detectability of IPATS-BLV and combined PCR-RFLP-sequencing.

|  |  | Combined PCR-RFLP-sequencing |  |  |  |
| --- | --- | --- | --- | --- | --- |
|  |  | *DRB3\*016:01* | *DRB3\*009:02* | *DRB3\*016:01/DRB3\*009:02* | Other alleles |
| IPATS-BLV | *DRB3\*016:01*/other allele<sup>a</sup> | 2 | 0 | 0 | 0 |
|  | *DRB3\*009:02*/other allele<sup>b</sup> | 0 | 9 | 0 | 0 |
|  | *DRB3\*016:01/DRB3\*009:02* | 0 | 0 | 5 | 0 |
|  | Other alleles | 0 | 0 | 0 | 42 |

<sup>a</sup>except *DRB3\*009:02*
<sup>b</sup>except *DRB3\*016:01*

### BLV infection diagnostic performance of IPATS-BLV is comparable with that of other diagnostic methods

We first evaluated the BLV infection diagnostic performance of IPATS-BLV by comparing it with that of the anti-gp51 antibody ELISA test. We performed both the ELISA test and IPATS-BLV for 65 samples with an unknown infectious status. We qualitatively compared the ELISA-positive/negative results versus the IPATS-BLV-positive/negative results. As shown in Table 2, 27 samples were identified as BLV-positive and 33 samples as BLV-negative by both assays. One sample was identified as BLV-positive by IPATS-BLV but as BLV-negative by ELISA; this discrepancy could result from a sample taken during the initial phase of BLV infection. Four samples were identified as BLV-negative by IPATS-BLV but as BLV-positive by ELISA. This result might indicate that these cattle were capable of suppressing an increase in the BLV PVL. Among these cattle, one was identified as carrying the *DRB3*009:02* allele. The kappa value between the IPATS-BLV and ELISA was 0.8452 (SE: ±0.1235).

**Table 2.**
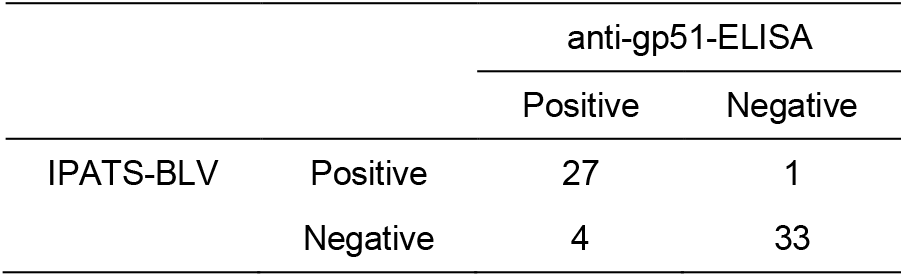
Comparison of the BLV detectability of IPATS-BLV and ELISA.

|  |  | anti-gp51-ELISA |  |
| --- | --- | --- | --- |
|  |  | Positive | Negative |
| IPATS-BLV | Positive | 27 | 1 |
|  | Negative | 4 | 33 |

Next, we evaluated the accuracy of the measurement of the percentage of BLV-infected cells by IPATS-BLV via a comparison with qPCR. We found a strong correlation (Pearson’s coefficient R = 0.9858, *p* < 1×10 ^15^) between these two assays, based on the measurement of 40 samples with variation in their percentage of BLV-infected cells (Figure 3). Finally, we determined the limit of detection (LOD) of the percentage of BLV-infected cells in IPATS-BLV using DNA extracted from serially diluted whole blood of BLV-infected cattle. IPATS-BLV could detect BLV provirus from cattle in which 1.50×10^-1^ percent of cells were infected with BLV, which is comparable to the LOD of commercial qPCR for BLV provirus (Table 3).

**Figure 3.**
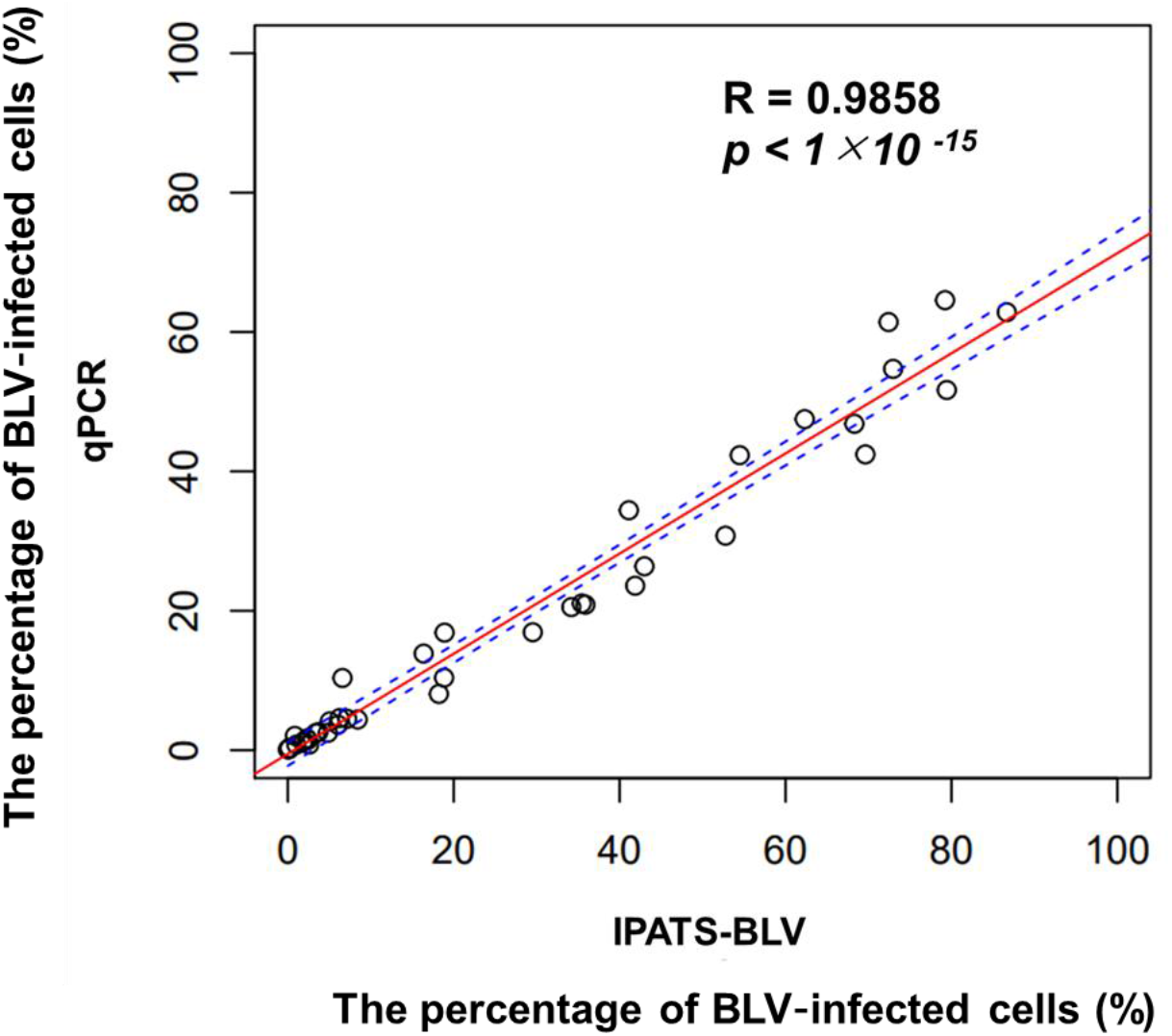
Correlation analysis of the measurement of the percentage of BLV-infected cells between IPATS-BLV and qPCR. The red line and blue dotted line indicate the linear model and 95% CI, respectively.

**Table 3.** Comparison of the BLV LOD between qPCR and IPATS-BLV.

| Percentage of infected cells (%) |  | qPCR | IPATS-BLV |  |
| --- | --- | --- | --- | --- |
|  |  | Ct | The No. of positive droplet | CNV2<sup>a</sup> |
| 1.50 | Fraction 1 | 34.71 | 39 | 0.014444 |
|  | Fraction 2 | 34.32 | 37 | 0.016897 |
|  | Fraction 3 | 34.68 | 46 | 0.018678 |
| 1.50×10<sup>-1</sup> | Fraction 1 | 38.08 | 5 | 0.001672 |
|  | Fraction 2 | 38.3 | 3 | 0.001059 |
|  | Fraction 3 | 39.52 | 6 | 0.002066 |
| 1.50×10<sup>-2</sup> | Fraction 1 | Undetected | 1 | 0.000326 |
|  | Fraction 2 | 40.65 | 0 | NA<sup>b</sup> |
|  | Fraction 3 | Undetected | 0 | NA |
| 1.50×10<sup>-3</sup> | Fraction 1 | Undetected | 0 | NA |
|  | Fraction 2 | Undetected | 0 | NA |
|  | Fraction 3 | Undetected | 0 | NA |

<sup>a</sup>BLV copy number per two RPP30 copies
<sup>b</sup>NA: Not available

### Survey for the percentage of DRB3*016:01- and DRB3*009:02-carrying cattle, and impact of these alleles on the percentage of BLV-infected cells

A field survey of the percentage of *DRB3*016:01* or *DRB3*009:02*-carrying cattle and the impact of these alleles on the BLV PVL was carried out in Miyazaki prefecture, Japan. First, we used an anti-gp51 ELISA to screen for BLV-infected cattle. Among 4,603 asymptomatic Japanese Black cattle from 1,394 farms, 353 cattle (7.7%) from 164 farms were identified as BLV-positive by ELISA (“ELIZA-positive”). We then performed IPATS-BLV on samples from the 353 ELISA-positive cattle; 200 cattle (56.7%) and 24 cattle (6.8%) were found to carry *DRB3*016:01* and *DRB3*009:02,* respectively. Prior to performing a comparison of the percentage of BLV-infected cells, we classified these cattle into the following five groups: *DRB3*016:01/*009:02* heterozygous (n = 8), *DRB3*009:02/Other* allele heterozygous (n = 16), *DRB3*016:01/*016:01* homozygous (n = 37), *DRB3*016:01*/Other allele heterozygous (n = 155), and Other alleles (n = 137) (Figure 4). The 37 *DRB3*016:01/*016:01* homozygous cattle showed an average *DRB3*016:01* ratio of 0.9930 (SE: ±0.014965). Cattle with a *DRB3*009:02* allele had a significantly lower percentage of BLV-infected cells compared with the in the other groups, even when the cattle were heterozygous for the BLV-susceptible *DRB3*016:01* allele. Although cattle with a *DRB3*016:01* allele had a statistically significantly higher percentage of BLV-infected cells compared with other allele-carrying cattle, their PVLs varied widely.

**Figure 4.**
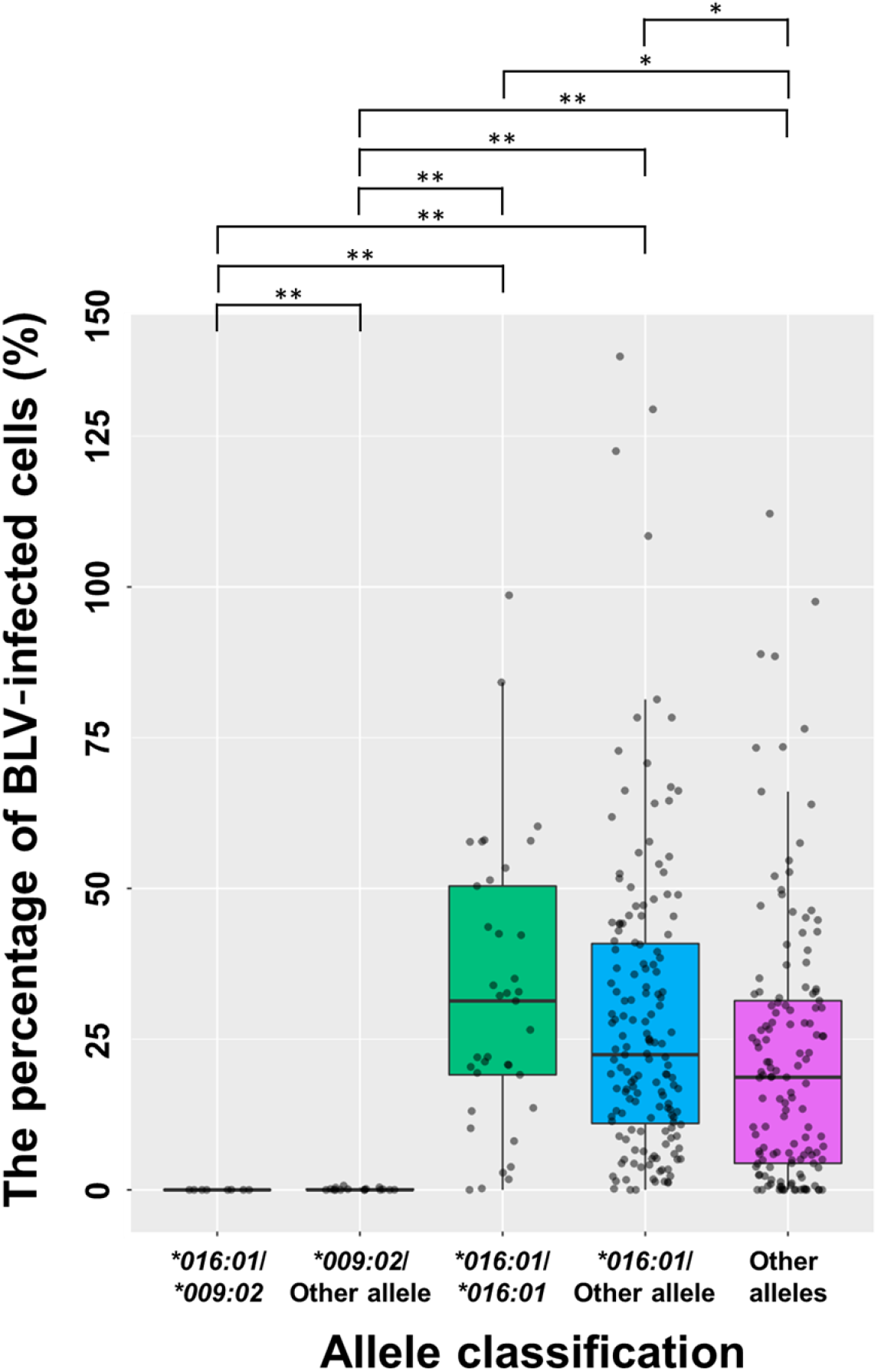
Comparison of the percentage of BLV-infected cells by allele classification. A box-and-whisker plot is shown. Box: 25^th^–75^th^ percentile of the range of the percentage of BLV-infected cells. Intermediate line in the box: Median. Dot: Each sample. \**p* < 0.5; \*\**p* < 0.0001

## Discussion

This study successfully developed a simple and relatively speedy test for both host genetic susceptibility and pathogen quantity, which will provide a deeper understanding of infection in individual patients and guide their appropriate treatment. An important usage of this platform is a risk analysis of the transmissibility of infected animals for veterinary science. We developed the IPATS-BLV method to identify BLV-susceptible super spreaders and BLV-resistant elite controllers more easily and rapidly. This test provides an absolute DNA quantification of the BLV *pol* gene, BLV-susceptible *DRB3*016:01* allele, BLV-resistant *DRB3*009:02* allele, and RPP30 by using a fourplex ddPCR. IPATS-BLV was demonstrated to accurately measure the percentage of BLV-infected cells and provide highly sensitive and specific allele typing that discriminates between homozygous and heterozygous carriers, all in a single-well reaction. We found that cattle carrying the BLV-resistant *DRB3*009:02* allele had a strong ability to maintain the PVL of BLV at a low or undetectable level. In contrast, *DRB3*016:01*-carrying cattle were found to have a relatively higher percentage of BLV-infected cells when compared with other allele-carrying cattle.

Here, we demonstrated the allelic impact of the previously identified BLV-susceptibility *DRB3*016:01* allele and BLV-resistant *DRB3*009:02* allele on the BLV PVL, as shown in Figure 4. *DRB3*009:02*-carrying cattle had a low/undetectable level of BLV provirus, even when their other allele was the BLV-susceptible *DRB3*016:01* allele. This result is supported by previous studies, indicating a strong association between *DRB3*009:02* and a low/undetectable PVL of BLV under the consideration of allele heterozygosity (El Daous et al., 2021; Lo et al., 2021). However, not all *DRB3*009:02-carrying* cattle are BLV resistant (Farias et al., 2017). It seems that BLV resistance is determined by not only the *DRB3* allelic effect but also other factors, such as species and climate. One advantage of IPATS-BLV is that it identifies BLV-resistant elite controllers on the basis of both *DRB3*009:02* and undetectable BLV provirus. Notably, *DRB3*016:01-carrying* cattle had a significantly higher PVL of BLV when compared with cattle with other alleles. This is supported by previous study, indicating that the percentage of BLV HPL cattle was higher among the group of *DRB3*016:01* -carrying cattle (Miyasaka et al., 2013). However, our results also suggest that the PVL of *DRB3*016:01-carrying* cattle varies widely. As BLV susceptibility is a relative property at the population level, BLV-susceptible *DRB3*016:01* does not have sufficient power to strongly associate with BLV HPL, unlike the strong association between *DRB3*009:02* and low/undetectable BLV PVL. A population of BLV-susceptible allele-carrying cattle with low or undetectable BLV provirus was previously found (Nakatsuchi et al., 2022). The association between *DRB3*016:01* and BLV HPL seems to be limited in particular situations. When the property of HPL is derived from genetic susceptibility, BLV-susceptible HPL cattle are considered to maintain a HPL and transmit BLV to others over a long span. BLV-susceptible allele (*DRB3*015:01*)-carrying Holstein cattle with a HPL continued to have a HPL over a long observation period (Bai et al., 2021). We recommend prioritizing the isolation of cattle with both *DRB3*016:01* and a HPL of BLV among BLV-infected cattle.

The simultaneous detection of pathogens and host biomarkers contributes to strengthening the control of livestock infectious diseases. Because there are presently no vaccines or effective treatments for BLV infection, prevention is only available countermeasure. BLV was previously eliminated in some countries in Europe via the identification and stamping out of infected animals and the restriction of between-farm cattle movement from infected farms (Maresca et al., 2015; Nuotio et al., 2003). As the BLV PVL varies by individual, depending on the virus–host interaction and other factors, not all infected cattle pose a risk of transmitting BLV to other cattle. Recently, BLV control on the basis of the PVL has been implemented under the presumption that cattle with a LPL have low or no risk of BLV transmission (Marcela. A. Juliarena et al., 2016; Mekata et al., 2015; Ruggiero et al., 2019). In addition to viral factors, host factors such as the *DRB3* haplotype have also received focus as an indicator of BLV disease susceptibility (Takeshima & Aida, 2006). Several studies identified some *DRB3* alleles as being associated with a LPL, including the strongly resistant *DRB3*009:02* (Carignano et al., 2017; El Daous et al., 2021; Hayashi et al., 2017; M. A. Juliarena et al., 2008; Lo et al., 2021; Miyasaka et al., 2013; Takeshima et al., 2019). The identification of BLV elite controllers will be useful in disrupting the chain of BLV transmission (Marcela. A. Juliarena et al., 2016). Despite of the benefit of herd management conducted on the basis of both PVL and *DRB3* haplotype, it is too time-consuming to implement if PVL measurement and allele typing need to be performed independently. Our newly developed method allows these data to be obtained more easily and rapidly and could be further applied to a high-throughput diagnosis. The power of IPATS-BLV opens a new avenue of BLV control by permitting the consideration of both PVL and genetic susceptibility. Disease control using resistant animals has an aspect of providing assurance for food safety. Because of the genetic variation in susceptibility to infectious diseases among species, derived from co-evaluation with pathogens (Duxbury et al., 2019; O’Brien & Evermann, 1988), a population of livestock possessing the power of disease resistance should exist latently everywhere. As selective breeding is an applied use of natural resources, there is no need to evaluate its adverse health effects to humans, unlike products of genome engineering. In the case of genetically modified crops, commercialization requires 13 years from project development and 35.01 million US$ for the cost of regulatory safety assessment and of securing global registration and authorizations. Notably, it takes five to seven years to perform the safety evaluations and obtain regulatory (Kumar et al., 2020). Ethical problems are also unavoidable when applying genome engineering to animals. Taken together, despite the advantage of the customizability of genome engineering for livestock, there is a bottleneck for implementing this approach. Genetic selection, which is already performed largely in marine (D’Agaro et al., 2021), forest (Lebedev et al., 2020), and livestock agriculture (Hayes et al., 2013), is a feasible alternative to genome engineering. This technique is ready to use when the equipment for selective breeding and diagnostics is available.

Consideration of both host biomarkers and pathogen levels has the potential for improving decision-making regarding the treatment and prevention of infectious diseases by providing a deeper understanding of individual infection. For example, septic shock outcome can be successfully predicted by merging information about the quantity of cytokines and bacteria in a patient (Abasiyanik et al., 2020). Regarding the current outbreak of severe acute respiratory syndrome coronavirus 2 (SARS-CoV-2) infection, researchers are discussing that HLA typing with viral diagnosis could improve the assessment of disease severity and allow high-risk individuals to be prioritized for vaccination (Nguyen et al., 2020). Such concepts contribute to improving preventive veterinary medicine by supporting appropriate herd management. Even when there are effective treatments and vaccinations for some threatening infectious diseases, some countries have a distribution bottleneck for these pharmacologic compounds owing to complex matters including supply chain and equipment (Acosta et al., 2019). Managing animals according to their current and future risk of disease transmissibility results in the best usage of available bioresources to suppress the damage from infectious diseases. Therefore, we expect the power of improved diagnostics to contribute to sustainable production from livestock in the future.

Some limitations of this study must be discussed. First, *DRB3*009*02*-carrying cattle with undetectable provirus could be either a BLV elite controller or an uninfected animal. We recommend the use of IPATS-BLV in combination with an antibody detection method, such as an ELISA. Second, *DRB3*009:02*-carrying cattle can have detectable BLV provirus in the initial phase of BLV infection (Forletti et al., 2020). Thus, the determination of BLV-resistant cattle should be conducted by testing the PVL several times.

In conclusion, IPATS is an easy and rapid platform with which to measure host biomarkers and pathogen levels. It provides strengthened diagnostics that consider both the disease susceptibility of the host and the actual disease severity/transmissibility. Such an approach has the potential to become a key tool for next-generation human and veterinary medicine.

## Materials and Methods

### IPATS-BLV assay design

We designed a fourplex ddPCR based on BLV proviral DNA, *DRB3*009:02, DRB3*016:01,* and RPP30-TaqMan Assay (Figure 1A–1F). To address the limited number of channels in our commercial ddPCR system (e.g., QX200 Droplet Digital PCR system, Bio-Rad, Hercules, USA), we modulated the amplicon length and primer/probe concentration in the reaction mixture to enable the separation of different targets within the same color (Levy et al., 2021; Miotke et al., 2014). We set the FAM_low, FAM_high, HEX_low, and HEX_high channels to *DRB3*016:01,* BLV *pol* gene, *DRB3*009:02,* and RPP30, respectively.

### Primer/Probe

We obtained 382 sequences of *DRB3.2* alleles from the IPD-MHC database (EBML-EBI, 2021). For *DRB3*009:02,* we designed allele-specific primers and probe via minor modification of a previously developed *DRB3*009:02*-TaqMan assay^33^. To discriminate *DRB3*016:01*, we designed a *DRB3*016:01*-specific forward primer and probe. The *DRB3*016:01*-TaqMan assay shares the reverse primer for *DRB3*009:02.* One concern of this design was potential nonspecific reactions between the *DRB3*009:02*-primer/probe and *DRB3*009:02*-primer/probe. Thus, we recruited LNA primers to suppress the undesired amplification of untargeted alleles. To detect wild strains of BLV with sequence diversity, we designed primers and probe targeting a conserved region in the *pol* gene, as identified from a database of aligned sequences for 82 reported strains (Table S4). This database includes 72 strains of BLV genotype 1 (G1), which is currently dominant worldwide, one strain of G2, one strain of G4, three strains of G6, four strains of G9, and one strain of G10. The primers and probe target a position in the 3’ terminal end of the *pol* gene (Figure S2), that is conserved except for an acceptable mismatch at the 5’ side of the forward primer in the par91 strain (Acc. No. LC080658.1). We added primers and probe for RPP30 into the reaction for housekeeping purposes. Table S1 indicates the sequences of the primers/probes. We purchased all these primers and probes, except for the LNA primers, from Eurofins Genomics (Tokyo, Japan). We purchased the LNA primers from QIAGEN (Hilden, Germany).

### IPATS-BLV

We finalized the IPATS-BLV reaction in a 22-μl reaction mixture containing 14 μl of 2× ddPCR Supermix for Probes (Bio-Rad, #1863023), 909 nM of primers except the RPP30 primers (*DRB3*016:01*-forward, *DRB3*009:02*-forward, *DRB3*009:02*-reverse, BLV *pol* 4527-forward, and BLV *pol* 4638-reverse), 455 nM of RPP30-forward and reverse primers, 68 nM of FAM-labeled *DRB3*016:01-probe,* 182 nM of HEX-labeled *DRB3*009:02*-probe, 295 nM of FAM-labeled BLV *pol* 4560-probe, 364 nM of HEX-labeled RPP30-probe, the sample adjusted to <35 ng, and the necessary volume of water to reach 22 μl (Table S2). We emulsified the reaction mixture using an automated droplet generator (#1864101JA, Bio-Rad) for partitioning into droplets in accordance with the manufacturer’s instructions. We performed PCR amplification according to the following amplification profile: 95 °C for 10 min; 60 cycles of 94 °C for 30 s and 58 °C for 2 min; 98 °C for 10 min. The FAM and HEX fluorescence magnitude of each droplet were read using a QX200™ Droplet Reader (#1864003JA, Bio-Rad). The number of droplets in each cluster was quantified by automatically/manually setting the appropriate fluorescence amplitude thresholds using QX Manager Software Standard Edition, Version 1.2 (Bio-Rad). We calculated the percentage of BLV-infected cells using below equation.

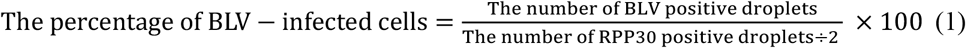

By calculating the ratio of the number of allele-positive droplets to the number of housekeeping-positive droplets using the below equation, we successfully discriminated whether cattle carry homozygous or heterozygous target alleles.

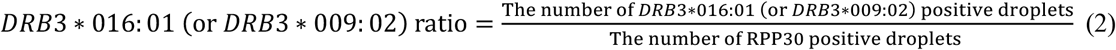

Ratios of approximately 1 and 0.5 indicate homozygosity and heterozygosity of an allele, respectively.

### Sensitivity and specificity of DRB3*009:02 and DRB3*016:01 genotyping

To determine the accuracy of *DRB3*009:02* and *DRB3*016:01* genotyping in IPATS-BLV, we genotyped 58 bovine genomic DNAs with varied *DRB3* alleles by IPATS-BLV. These samples included 21 *DRB3* alleles (Table S3), according to the results of *DRB3* allele determination using combined PCR-RFLP-sequencing methods (El Daous et al., 2021; Notsu et al., 2022; Van Eijk et al., 1992). The agreement of *DRB3.2* allele typing between combined PCR-RFLP-sequencing and IPATS-BLV was judged by calculating the diagnostic sensitivity and specificity.

### Agreement with commercial ELISA

We judged the agreement of qualitative detectability of BLV-infected cattle of IPATS-BLV with a commercial anti-gp51 antibody ELISA kit (#No cat. number, Nippon gene, Tokyo, Japan). In the experiment, we used 65 bovine blood samples of unknown BLV infectious status. We isolated plasma by centrifuging the samples for 10 min at 1000 ×*g*. The ELISA test was performed in accordance with the manufacturer’s instructions. We extracted genomic DNA from whole blood using a Wizard^®^ Genomic DNA Purification Kit (#A1120, Promega Corp., Madison, USA) and then performed IPATS-BLV. We defined samples as ELIS-Apositive if their value was higher than the cut-off S/P value and as IPATS-BLV-positive if more than one BLV-positive droplet was detected in the amplitude. We evaluated the consensus of ELISA-positive/negative versus IPATS-BLV-positive/negative by calculating a kappa value using software in epitools (Sergeant, 2018).

### Quantitativity of the percentage of BLV-infected cells

For the accuracy of the quantification of the percentage of infected cells in IPATS-BLV, we determined the correlation of measurement with a commercial qPCR kit (#RC202A, TaKaRa, Shiga, Japan). The commercial qPCR kit targeted he BLV *pol* gene and RPPH1 for housekeeping. We extracted genomic DNA samples from the whole blood of cattle using MagDEA Dx SV reagent (#E1300, Precision System Science, Chiba, Japan) with an automated nucleic acid extraction system (magLEAD 12gC, #A1120, Precision System Science) in accordance with the manufacturer’s instructions. Next, we performed qPCR in accordance with the manufacturer’s instructions. We selected 40 samples satisfying the variation of the percentage of infected cells and performed IPATS-BLV on these samples. The strength of correlation between qPCR and IPATS-BLV was determined using Pearson’s coefficient, calculated using R software v. 3. 6. 2 (R Development Core Team, 2019).

### LOD of BLV detection

To determine the LOD of BLV detection in IPATS-BLV, we tested DNA samples extracted from a serial dilution series of whole blood from BLV-infected cattle. This animal carried 1.5% of BLV-infected cells (as confirmed using qPCR). We serially diluted the whole blood of this animal 10 times using whole blood from a BLV-uninfected animal. We confirmed the “uninfected” status of these cattle by both the absence of provirus in a qPCR assay and the absence of anti-BLV gp51 antibody in an ELISA. We extracted genomic DNA from three fractions of each dilution using magLEAD 12gC. We performed both IPATS-BLV and qPCR to compare the LOD. In both assays, the sample DNA input in the reaction mixture was 20 ng.

### Field survey

We performed a field survey for the percentage of *DRB3*016:01-* or *DRB3*009:02*-carrying cattle and the impact of these alleles on the BLV PVL. We targeted asymptomatic Japanese Black cattle in Miyazaki prefecture, Japan. Whole blood samples were collected from 4,603 cattle over 1,394 farms by veterinarians and sent to University of Miyazaki. These samples were collected from May 2020 to July 2022. Anti-BLV gp51 antibody ELISA tests were performed immediately to screen for BLV-infected cattle. We stored the whole blood of ELISA-positive samples at −20 °C until their use in further analysis. We extracted the genomic DNA of ELISA-positive cattle using either the magLEAD 12gC or a MagMAX™ CORE Nucleic Acid Purification Kit (Thermo Fisher Scientific Inc., Waltham, USA) with an automated nucleic acid extraction system (KingFisher Duo Prime; Thermo Fisher Scientific Inc.). We performed IPATS-BLV for *DRB3*016:01*, *DRB3*009:02*, and BLV PVL. We classified these samples into the following five groups: *DRB3*016:01*/*DRB3*009:02*, *DRB3*009:02*/other allele, *DRB3*016:01*/*DRB3*016:01* (*DRB3*016:01* homozygous), *DRB3*016:01* /other allele, and other alleles groups prior to a comparison of the percentage of BLV-infected cells between groups. We used a pairwise Wilcoxon rank sum test with Bonferroni’s modification for determining the significance of differences between each group using R software. Differences with a *p*-value of <0.05 were judged as statistically significant.

## Acknowledgements

Research reported in this publication was supported by Grant-in-Aid for JSPS Fellows Grant Number JP21J23396 (K.N) and JSPS KAKENHI Grant Number JP20K06413 (S.S). We thank Toshie Iwanaga and Rika Nohara at University of Miyazaki for data collection assistance and Drs. Chika Ryu and Yuchi Ushitani at JA Miyazaki for coordinating the sample collections. We also thank Katie Oakley, PhD, from Edanz (https://jp.edanz.com/ac) for editing a draft of this manuscript.

## Author contributions

K.N. and S.S. conceived of this study and acquired funding; S.S. supervised the project; K.N. designed the IPATS-BLV protocol and performed all IPATS-BLV experiments; K.N., H.E.D., and S.M. performed the *DRB3* allele typing by using combined PCR-RFLP-sequencing; K.N. performed the BLV anti-gp51-ELISA test; K.N. and X.W. performed the BLV qPCR; and K.N. prepared the manuscript and figures. All authors read, revised, and approved the manuscript.

## Competing interests statement

The authors declare that they have no conflict of interest.

## Notes

### Competing Interest Statement

The authors have declared no competing interest.

