## Supplementary Information for "Identifying Pathogen and Allele Type Simultaneously (IPATS) in a single well using droplet digital PCR"

#### Contents

Figure S1: Cluster pattern in 1D amplitude of IPATS-BLV. Page 2.

Figure S2: Mutation mapping in BLV *pol* gene and location of primers and probe. Page 3.

Table S1: IPATS-BLV primers and probes. Page 4.

Table S2: Composition of IPATS-BLV reaction mixture. Page 4.

Table S3: Genotyping of various alleles using IPATS-BLV. Page 5.

Table S4: List of aligned BLV strains to determine conserved region. Page 6.

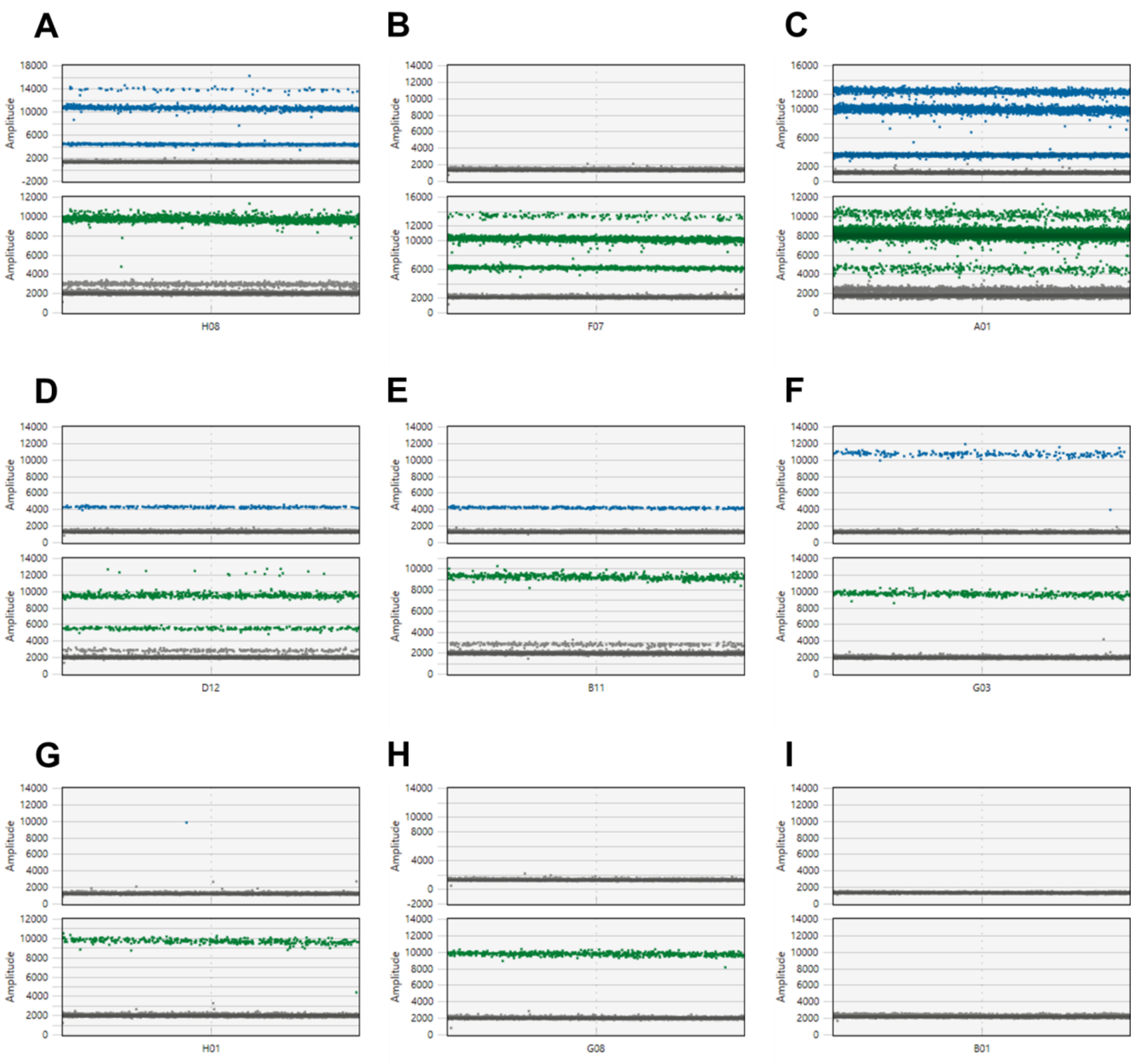


**Figure. S1. Cluster pattern in 1D amplitude of IPATS-BLV.**
Above and below indicates FAM and HEX amplitude, respectively. (A) BLV-susceptible super spreader. (B) BLV-resistant elite controller. (C) Mixed population of *DRB3*016:01/*015:01*-carrying cattle with a HPL of BLV and *DRB3*009:02/*015:01*-carrying cattle (presumably *DRB3*009:02/*016:01* heterozygous with detectable BLV provirus). (D) *DRB3*009:02/*016:01* heterozygous cattle with undetectable BLV provirus. (E) *DRB3*016:01*-carrying cattle with undetectable BLV provirus. (F) Other allele-carrying cattle with a HPL of BLV. (G) Other allele-carrying cattle with a LPL of BLV. (H) Other allele-carrying cattle with undetectable BLV provirus. (I) Water.

#### > pvAN003 strain, Acc. No. AP018024.1 pol gene 2343-4800nt

```
CTAGAACGCCTCCAGGCCCTTCAAGACCTGGTCCATCGCTCTCTGGAGGCAGGTTATATCTCCCCC
TGGGACGGGCCAGGCAATAATCCAGTCTTCCCGGTACGGAAACCAAATGGCGCCTGGAGGTTTGT
GCATGACCTACGAGCTACAAATGCTCTTACAAAGCCCATTCCGGCACTCTCCCCCGGACCGCCAG
ACCTTACCGCTATCCCTACACACCTTCCACATATCATTTGCCTAGATCTCAAAGACGCCTTCTTCCA
GATTCCAGTCGAAGACCGCTTCCGCTCCTACTTTGCTTTTACCCTCCCTACCCCCGGGGGACTCCA
ACCTCATAGACGCTTTGCCTGGCGGGTCCTACCTCAAGGCTTCATTAACAGCCCAGCTCTTTTCGA
ACGAGCACTACAGGAACCTCTTCGCCGAGTTTCCGCCGCCTTTTCCCAGTCTCTTCTGGTGTCCTAT
ATGGACGATATCCTTATCGCTTCGCCTACAGAAGAACAGCGGTCACAATGTTATCAAGCCCTGGCT
GCCCGCCTCCGGGACCTAGGGTTTCAGGTGGCATCCGAAAAGACTCGCCAGACGCCTTCGCCCGT
CCCCTTCTTGGGACAAATGGTCCATGAGCAGATTGTCACCTACCAGTCCCTACCTACCTTGCAGAT
CTCATCCCCAATTTCTCTCCACCAATTACAGGCGGTCTTAGGAGACCTCCAATGGGTCTCTAGGGG
CACACCCACTACCCGCCGGCCCCTGCAACTTCTCTACTCTTCCCTTAAAGGCATCGATGACCCTAG
GGCCATCATCCAGCTTTCCCCGGAACAGCTGCAAGGCATTGCAGAGCTTCGACAAGCCCTGTCCC
ATAACGCAAGATCTAGATACAACGAGCAAGAACCCCTGCTAGCCTACGTACACCTAACCCGGGCG
GGGTCCACCCTGGTACTCTTCCAAAAGGGCGCTCAATTTCCCCTGGCCTACTTCCAGACCCCCTTG
ACTGACAACCAAGCCTCACCTTGGGGCCTCCTTCTCCTGCTGGGATGCCAATACCTGCAGACTCAG
GCCTTAAGCTCGTATGCCAAGCCCATACTTAAATATTATCACAATCTTCCTAAAACTTCTCTAGACA
ATTGGATTCAATCATCTGAGGACCCTCGAGTCCAGGAGTTGCTGCAATTGTGGCCCCAGATTTCCT
CTCAGGGAATACAGCCCCCGGGCCCTTGGAAGACCTTAATCACCAGGGCAGAGGTTTTTTTGACG
CCCCAGTTCTCCCCTGATCCGATTCCTGCGGCCCTTTGCCTCTTTAGTGACGGGGCTAAAGGACGA
GGAGCATATTGCTTGTGGAAAGACCACCTTTTGGACTTTCAGGCCGTTCCGGCTCCAGAATCCGCT
CAAAAGGGAGAACTGGCAGGTCTCTTGGCGGGCTTAGCAGCCGCCCCGCCTGAACCTGTAAATAT
ATGGGTAGATTCCAAATACCTGTACTCTTTGCTCAGAACCCTAGTTCTGGGGGCTTGGCTTCAACCT
GACCCCGTACCCTCCTACGCCCTCCTATATAAAAGCCTCCTCCGACATCCAGCAATCTTTGTTGGC
CATGTCCGGAGCCACTCTTCAGCATCCCACCCTATTGCTTCCCTGAACAATTATGTAGATCAACTG
CTTCCCTTAGAAACTCCAGAGCAATGGCATAAGCTCACCCACTGCAACTCTCGGGCCTTGTCTCGA
TGGCCGAACCCACGTATCTCTGCCTGGGACCCCCGTTCCCCCGCTACGCTGTGTGAAACCTGTCAA
AAGCTTAATCCAACTGGAGGAGGAAAGATGCGAACTATTCAGAGAGGGTGGGCCCCGAACCATAT
TTGGCAGGCCGATATAACCCATTATAAATACAAACAGTTCACCTACGCTCTGCATGTGTTTGTAGAT
ACTTACTCTGGAGCTACTCATGCCTCGGCGAAGCGTGGGCTCACCACTCAAATGACCATTGAGGG
CCTTCTTGAGGCCATAGTGCATCTGGGTCGTCCAAAAAAGCTAAACACTGACCAAGGTGCAAACTA
CACCTCCAAAACCTTTGTCAGGTTTTGCCAGCAGTTCGGAGTTTCCCTTTCTCATCATGTTCCCTAC
AACCCCACAAGTTCGGGGTTAGTAGAACGGACAAATGGACTGCTCAAACTTCTTCTATCTAAATAT
CACCTAGACGAACCCCACCTTCCCATGACTCAGGCCCTTGCTCGAGCCCTCTGGACTCACAATCA
GATTAACCTCCTACCAATTCTAAAGACCAGATGGGAGCTACACCATTCACCCCCACTTGCTGTCAT
TTCAGAGGGCGGAGAAACACCCAAGGGCTCTGATAAACTCTTTTTGTACAAGCTCCCCGGGCAAA
ACAATCGTCGGTGGCTAGGACCACTCCCGGCCCTAGTCGAAGCCTCGGGAGGCGCTCTCCTGGCT
ACTGACCCCCCCGTGTGGGTTC
```

**Figure. S2. Mutation mapping in BLV *pol* gene and location of primers and probe.**

Nucleotides at least 1 strain had mismatch are red-colored in a sequence of reference strain (pvAN003, Acc. No. AP018024.1). Primers and probe position are indicated by blue and pink blanket, respectively.

#### Table S1. IPATS-BLV primers and probes.

| Primer/Probe | Target | Sequence 5' to 3' | Acc. No. of reference | Position |
|---|---|---|---|---|
| *DRB3*016:01*-forward | *DRB3*016:01* | TTCGTGCGCTTCGA+T[a] | AB610127.1 | 107-121 |
| *DRB3*016:01*-probe | *DRB3*016:01* | FAM-AGGACTTCCTGGAGGAGAA-MGB-Eclipse | AB610127.1 | 192-210 |
| *DRB3*009:02*-forward | *DRB3*009:02* | GTGCGGTTCCTGGA+G[a] | AB610142.1 | 68-82 |
| *DRB3*009:02*-probe | *DRB3*009:02* | HEX-AGATCCTGGAGGAGAGGC-MGB-Eclipse | AB610142.1 | 195-212 |
| *DRB3*009:02*-reverse | *DRB3*016:01* and *DRB3*009:02* | CGCTGCACAGTGAAACTCTCA | AB610142.1 | 256-276 |
| BLV *pol* 4527-forward | BLV proviral DNA | GAACCCCACCTTCCCATGAC | AP018024.1 | 4,527-4,546 |
| BLV *pol* 4560-probe | BLV proviral DNA | FAM-CGAGCCCTCTGGACTCACAATC-BHQ1 | AP018024.1 | 4,560-4,581 |
| BLV *pol* 4638-reverse | BLV proviral DNA | GCCCTCTGAAATGACAGCAAG | AP018024.1 | 4,638-4658 |
| RPP30-forward | RPP30 | TGTTTCTGTTGGTCTGGTGTCC | NC_037353.1 | 12,515,902-12,515,923 |
| RPP30-probe | RPP30 | HEX-CGGCTGACTCTGGGCTGAA-MGB-Eclipse | NC_037353.1 | 12,515,925-12,515,943 |
| RPP30-reverse | RPP30 | CGGCCTTCGCATCACTTTC | NC_037353.1 | 12,515,984-12,516,002 |

[a]+N indicates LNA.

#### Table S2. Composition of IPATS-BLV reaction mixture.

| Reagent/Primer/Probe/Sample | ×1 | Final concentration |
|---|---|---|
| ddPCR™ Supermix for Probes (No dUTP) | 14 µl | - |
| *DRB3*016:01*-forward (40 µM) | 0.5 µl | 909 nM |
| *DRB3*016:01*-probe (10 µM) | 0.15 µl | 68 nM |
| *DRB3*009:02*-forward (40 µM) | 0.5 µl | 909 nM |
| *DRB3*009:02*-probe *(10* µM*)* | 0.4 µl | 182 nM |
| *DRB3*009:02*-reverse (40 µM) | 0.5 µl | 909 nM |
| BLV *pol* 4527-forward (40 µM) | 0.5 µl | 909 nM |
| BLV *pol* 4560-probe (10 µM) | 0.65 µl | 295 nM |
| BLV *pol* 4638-reverse (40 µM) | 0.5 µl | 909 nM |
| RPP30-forward (20 µM) | 0.5 µl | 455 nM |
| RPP30-probe (10 µM) | 0.8 µl | 364 nM |
| RPP30-reverse (20 µM) | 0.5 µl | 455 nM |
| $H_2O$ | 0.5 µl | - |
| DNA Sample (< 35 ng in reaction mixture) | 2 µl | - |
| Total | 22 µl | - |

### Table S3. Genotyping of various alleles using IPATS-BLV.

| Sample ID | Allele 1 | Allele 2 | *DRB3*016:01* ratio | *DRB3*009:02* ratio | The percentage of BLV-infected cells |
|---|---|---|---|---|---|
| #1 | *005:02 | *014:01:01 | 0.00036007 | 0.01191342 | ND[a] |
| #2 | *010:01 | *015:01 | ND | 0.01070361 | ND |
| #3 | *001:01 | *011:01 | ND | 0.02420252 | ND |
| #4 | *006:01 | *010:01 | ND | 0.00344443 | ND |
| #5 | *014:01:01 | *027:03 | ND | 0.00237900 | 0.4758005 |
| #6 | *006:01 | *015:01 | ND | 0.02617585 | 49.794859 |
| #7 | *010:01 | *027:03 | 0.00112552 | 0.00487844 | 2.4779443 |
| #8 | *001:01 | *001:01 | ND | 0.01094380 | 15.252295 |
| #9 | *001:01 | *012:01 | 0.00041453 | 0.00580529 | 138.28632 |
| #10 | *010:01 | *012:01 | ND | 0.00446447 | 73.646647 |
| #11 | *015:01 | *027:03 | 0.00730837 | 0.00584656 | 103.99457 |
| #12 | *010:01 | *012:01 | ND | ND | 80.204463 |
| #13 | *001:01 | *027:03 | 0.00181486 | 0.00181486 | 4.7199491 |
| #14 | *001:01 | *010:01 | ND | ND | 11.779267 |
| #15 | *010:01 | *027:03 | ND | 0.00612561 | 37.153152 |
| #16 | *002:01 | *012:01 | 0.00582382 | 0.01164793 | 29.136074 |
| #17 | *001:01 | *027:03 | 0.00544986 | 0.00181654 | 79.942411 |
| #18 | *011:01 | *012:01 | ND | 0.00429758 | 74.969059 |
| #19 | *001:01 | *001:01 | ND | ND | 74.046493 |
| #20 | *001:01 | *012:01 | 0.00211347 | 0.00211347 | 98.168093 |
| #21 | *010:01 | *015:01 | ND | 0.00956051 | 41.497043 |
| #22 | *011:01 | *027:03 | 0.00335384 | 0.00670784 | 71.959674 |
| #23 | *011:01 | *015:01 | ND | 0.00681452 | 67.910761 |
| #24 | *010:01 | *027:03 | ND | ND | 88.39801 |
| #25 | *011:01 | *015:01 | 0.00630761 | 0.01892364 | 79.582584 |
| #26 | *012:01 | *027:03 | ND | 0.00606147 | 91.92639 |
| #27 | *010:01 | *015:01 | ND | 0.00388764 | 61.278611 |
| #28 | *010:01 | *011:01 | ND | ND | 98.273414 |
| #29 | *009:02 | *015:01 | ND | 0.41412233 | ND |
| #30 | *002:01 | *012:01 | ND | 0.00918637 | 0.166988431 |
| #31 | *002:01 | *012:01 | ND | ND | ND |
| #32 | *005:03 | *016:01 | 0.45545114 | 0.00368242 | 16.566636 |
| #33 | *002:01 | *012:01 | ND | 0.00955920 | 0.802813 |
| #34 | *002:01 | *012:01 | ND | 0.00701826 | 0.0519577 |
| #35 | *005:03 | *002:01 | 0.00038352 | 0.00838206 | 17.991567 |
| #36 | *005:02 | *012:01 | ND | 0.00662005 | 1.7654261 |
| #37 | *009:02 | *015:01 | ND | 0.46336997 | ND |
| #38 | *005:03 | *034:01 | 0.00177072 | 0.00431248 | ND |
| #39 | *009:02 | *015:01 | ND | 0.48082397 | ND |
| #40 | *009:02 | *015:01 | ND | 0.50310229 | ND |
| #41 | *007:01 | *009:02 | 0.00360128 | 0.48387558 | ND |
| #42 | *007:01 | *009:02 | 0.00161369 | 0.48879834 | ND |
| #43 | *009:02 | *016:01 | 0.47247611 | 0.46766509 | ND |
| #44 | *009:02 | *016:01 | 0.50402739 | 0.49878371 | ND |
| #45 | *009:02 | *016:01 | 0.46188088 | 0.49132784 | ND |
| #46 | *009:02 | *016:01 | 0.42381518 | 0.43059562 | ND |
| #47 | *009:02 | *016:01 | 0.43270428 | 0.42206102 | ND |
| #48 | *009:02 | *010:01 | 0.00031968 | 0.47509541 | ND |
| #49 | *009:02 | *010:01 | ND | 0.42521606 | ND |
| #50 | *001:01 | *018:01 | 0.00060168 | 0.00692286 | ND |
| #51 | *012:01 | *015:01 | ND | ND | ND |
| #52 | *044:01 | *045:01 | ND | ND | ND |
| #53 | *012:01 | *014:01:01 | 0.00027370 | 0.00821650 | ND |
| #54 | *008:01 | *045:01 | 0.00048788 | 0.00707893 | ND |
| #55 | *037:01 | *044:01 | 0.19204542 | 0.00829384 | 0.1381294 |
| #56 | *005:02 | *009:02 | 0.00004299 | 0.47631446 | ND |
| #57 | *018:01 | *028:01 | ND | 0.00143383 | ND |
| #58 | *015:01 | *016:01 | 0.50207961 | 0.00252116 | 101.9198895 |

[a]ND: No positive droplet was detected.

### Table S4. List of aligned BLV strains to determine conserved region.

| Name of strain | Genotype | Acc. No. | Name of strain | Genotype | Acc. No. |
|---|---|---|---|---|---|
| AB934282.1[†] | 1 | AB934282.1 | pvAJ013 | 1 | AP019577.1 |
| JOTK | 1 | AB987702.1 | pvAJ014 | 1 | AP019578.1 |
| pvAF060 | 1 | AP018006.1 | pvAJ015 | 1 | AP019579.1 |
| pvAF076 | 1 | AP018007.1 | pvAJ016 | 1 | AP019580.1 |
| pvAF193 | 1 | AP018008.1 | pvAJ017 | 1 | AP019581.1 |
| pvAF245 | 1 | AP018009.1 | pvAJ018 | 1 | AP019582.1 |
| pvAF266 | 1 | AP018010.1 | pvAJ019 | 1 | AP019583.1 |
| pvAF293 | 1 | AP018011.1 | pvAJ021 | 1 | AP019585.1 |
| pvAF438 | 1 | AP018012.1 | pvAJ022 | 1 | AP019586.1 |
| pvAF481 | 1 | AP018013.1 | pvAJ023 | 1 | AP019587.1 |
| pvAF513 | 1 | AP018014.1 | pvAJ024 | 1 | AP019588.1 |
| pvAF746 | 1 | AP018015.1 | pvAJ025 | 1 | AP019589.1 |
| pvAF784 | 1 | AP018016.1 | pvAJ026 | 1 | AP019590.1 |
| pvAF805 | 1 | AP018017.1 | pvAJ027 | 1 | AP019591.1 |
| pvAF902 | 1 | AP018018.1 | pvAJ028 | 1 | AP019592.1 |
| pvAF982 | 1 | AP018019.1 | pvAJ029 | 1 | AP019593.1 |
| pvAK001 | 1 | AP018020.1 | pvAJ030 | 1 | AP019594.1 |
| pvAK006 | 1 | AP018021.1 | pvAJ031 | 1 | AP019595.1 |
| pvAK007 | 1 | AP018022.1 | pvAJ032 | 1 | AP019596.1 |
| pvAK011 | 1 | AP018023.1 | pvAJ033 | 1 | AP019597.1 |
| pvAN003 (Reference in this study) | 1 | AP018024.1 | pvAJ034 | 1 | AP019598.1 |
| pvAN004 | 1 | AP018025.1 | LS3 | 1 | HE967303.1 |
| pvAN006 | 1 | AP018026.1 | K02120.1[a] | 1 | K02120.1 |
| pvAN008 | 1 | AP018027.1 | *469, deficient type | 1 | LC005616.1 |
| pvAN009 | 1 | AP018028.1 | par7 | 1 | LC080653.1 |
| pvAN011 | 1 | AP018029.1 | pvAF019 | 1 | LC164084.1 |
| pvAN013 | 1 | AP018030.1 | pvAF967 | 1 | LC164085.1 |
| pvAN014 | 1 | AP018031.1 | pvAN903 | 1 | LC164086.1 |
| pvAN015 | 1 | AP018032.1 | BLV_BL3.1 | 1 | LC436098.1 |
| pvAJ001 | 1 | AP019565.1 | IBK1705 | 1 | LC552969.1 |
| pvAJ002 | 1 | AP019566.1 | V50F13 | 1 | MH170027.1 |
| pvAJ003 | 1 | AP019567.1 | Arg41 | 2 | FJ914764.1 |
| pvAJ004 | 1 | AP019568.1 | AF033818.1[a] | 4 | AF033818.1 |
| pvAJ005 | 1 | AP019569.1 | par62 | 6 | LC080656.1 |
| pvAJ006 | 1 | AP019570.1 | par91 | 6 | LC080658.1 |
| pvAJ007 | 1 | AP019571.1 | CHI-DQ | 6 | MG800834.1 |
| pvAJ008 | 1 | AP019572.1 | mon28 | 9 | LC080662.1 |
| pvAJ009 | 1 | AP019573.1 | por2 | 9 | LC080664.1 |
| pvAJ010 | 1 | AP019574.1 | por14 | 9 | LC080665.1 |
| pvAJ011 | 1 | AP019575.1 | por57 | 9 | LC080670.1 |
| pvAJ012 | 1 | AP019576.1 | QH2 | 10 | MF580995.1 |

[a]If strain name is not specific, named by accession number
